# SINBAD: a flexible tool for single cell DNA methylation data

**DOI:** 10.1101/2021.10.23.465577

**Authors:** Yasin Uzun, Wenbao Yu, Changya Chen, Kai Tan

**Affiliations:** Center for Childhood Cancer Research, The Children’s Hospital of Philadelphia, Philadelphia, PA 19104, USA; Department of Biomedical and Health Informatics, The Children’s Hospital of Philadelphia, Philadelphia, PA 19104, USA; Department of Genetics, University of Pennsylvania, Philadelphia, PA 19104, USA; Department of Pediatrics, University of Pennsylvania, Philadelphia, PA 19104, USA; Penn Epigenetics Institute, University of Pennsylvania, Philadelphia, PA 19104, USA

**Keywords:** epigenetics, single cell, DNA methylation, bisulfite sequencing

## Abstract

DNA methylation is an epigenetic mark that has vital importance in both development and disease. Single cell bisulfite sequencing technologies enable profiling of the methylome at high resolution, providing the basis for dissecting the heterogeneity and dynamics of DNA methylation in complex tissues and over time. Despite the rapid increase in the number of experimental protocols for methylome sequencing, analytical tools designed specifically for single-cell data are lacking. We developed a computational tool, SINBAD, for efficient and standardized pre-processing, quality assessment and analysis of single cell methylation data. Starting from multiplexed sequencing reads, major analysis modules of SINBAD include preprocessing, read mapping, methylation quantification, multivariate analysis, and gene signature profiling. SINBAD provides a flexible platform to implement interoperable and robust processing of single-cell methylome data.

## Background

DNA methylation plays a critical role in development and disease. It has been extensively studied using bulk samples with microarray and next-generation sequencing technologies (Lister et al. 2009; Jones 2012; Greenberg and Bourc’his 2019; Michalak et al. 2019). Due to its simplicity and ability to determine methylation state at single-nucleotide resolution, bisulfite sequencing (BS-Seq) is the most common experimental method for profiling the DNA methylome. Recently, a number of protocols have been developed to map DNA methylome at genome-wide coverage and single-cell resolution, mostly based on bisulfite sequencing (Karemaker and Vermeulen 2018; Stuart and Satija 2019; Ahn et al. 2021). Collectively, these single-cell protocols have revolutionized our ability to determine epigenetic heterogeneity in complex tissues and over time.

Unlike bulk DNA methylation data, single-cell methylome data poses unique analysis challenges. First, quality control needs to be performed at both bulk sample and single-cell level in order to detect and exclude low-quality cells. Second, even with deep sequencing, single-cell methylome data is inherently sparse for individual cells and only a small fraction of all cytosines across the genome can be covered by the assay. This presents a formidable challenge for estimating the methylation levels of the annotated regions for individual cells, and subsequent clustering to identify cell populations in the samples. Another challenge is the much larger size of single-cell methylome data compared to bulk methylome data. Unless the analysis procedures are designed and executed efficiently, the computational overhead can easily become prohibitive. For these reasons, computational methods designed for bulk methylation data are ill suited for single-cell data.

The initial processing step of single-cell methylome data includes demultiplexing, read alignment, cell-level quality control and methylation call. The quality of this initial processing step has a dramatic effect on downstream analyses such as dimensionality reduction, clustering, and integration with other types of single-cell omics data. Existing tools for single-cell DNA methylation data analysis ignore the initial processing step and only perform downstream processing to a limited extent, where the data is already pre-processed, aligned, quality controlled, and the methylation matrix is available at the gene level (Wolf, Angerer, and Theis 2018). As a result, in most single-cell DNA methylation studies, custom scripts are used for data processing and analysis, hampering standardization and reproducibility. There are a number of large-scale single-cell atlasing projects, including the Human Tumor Atlas Network (Rozenblatt-Rosen et al. 2020), the Brain Initiative Cell Census Network (Liu et al. 2021), The Human BioMolecular Atlas Program (HuBMAP, HuBMAP Consortium 2019) and the Human Cell Atlas (Rozenblatt-Rosen et al. 2017) in which multiple modalities of single-cell data, including DNA methylation, are produced. Depositing data processed with standardized and interoperable pipelines that are reproducible, is an indispensable component of these atlasing efforts. An efficient and flexible computational tool is critically needed to address this critical need.

## Results

### Overview of SINBAD

We addressed the lack of tools for single-cell methylation data QC and analysis by developing a flexible toolbox named SINBAD (A pipeline for processing SINgle cell Bisulfite sequencing samples and Analysis of Data). It consists of five analysis modules (Fig. 1). The pre-processing module performs demultiplexing of barcodes and trimming of adaptor sequences. The mapping module performs read alignment and filtering of low-quality reads and cells. Using filtered and aligned reads, the methylation module performs methylation call and quantification of cytosine sites of pre-specified genomic regions and generates a region-by-cell matrix of methylation levels. Next, the dimensionality of the methylation matrix is reduced by the multivariate analysis module and cell populations are detected by clustering analysis. Finally, the gene signature profiling module identifies methylation signatures of distinct cell populations by marker activities and differential methylation analyses.

**Figure 1.**
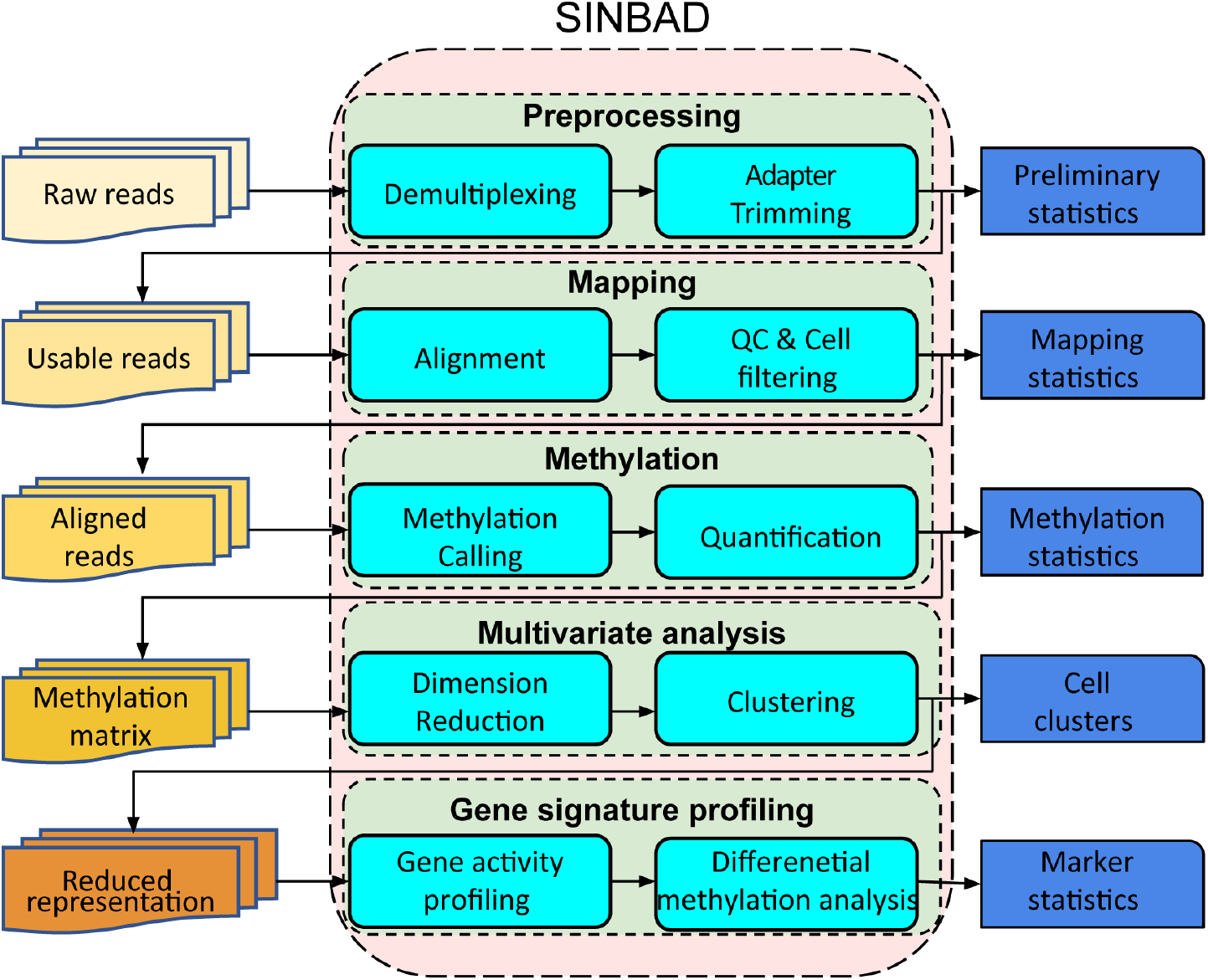
Overview of SINBAD. Left: Input and intermediate data. Middle, data processing and analysis modules. Right, Outputs generated by the software. QC, Quality Control.

**Figure 2.**
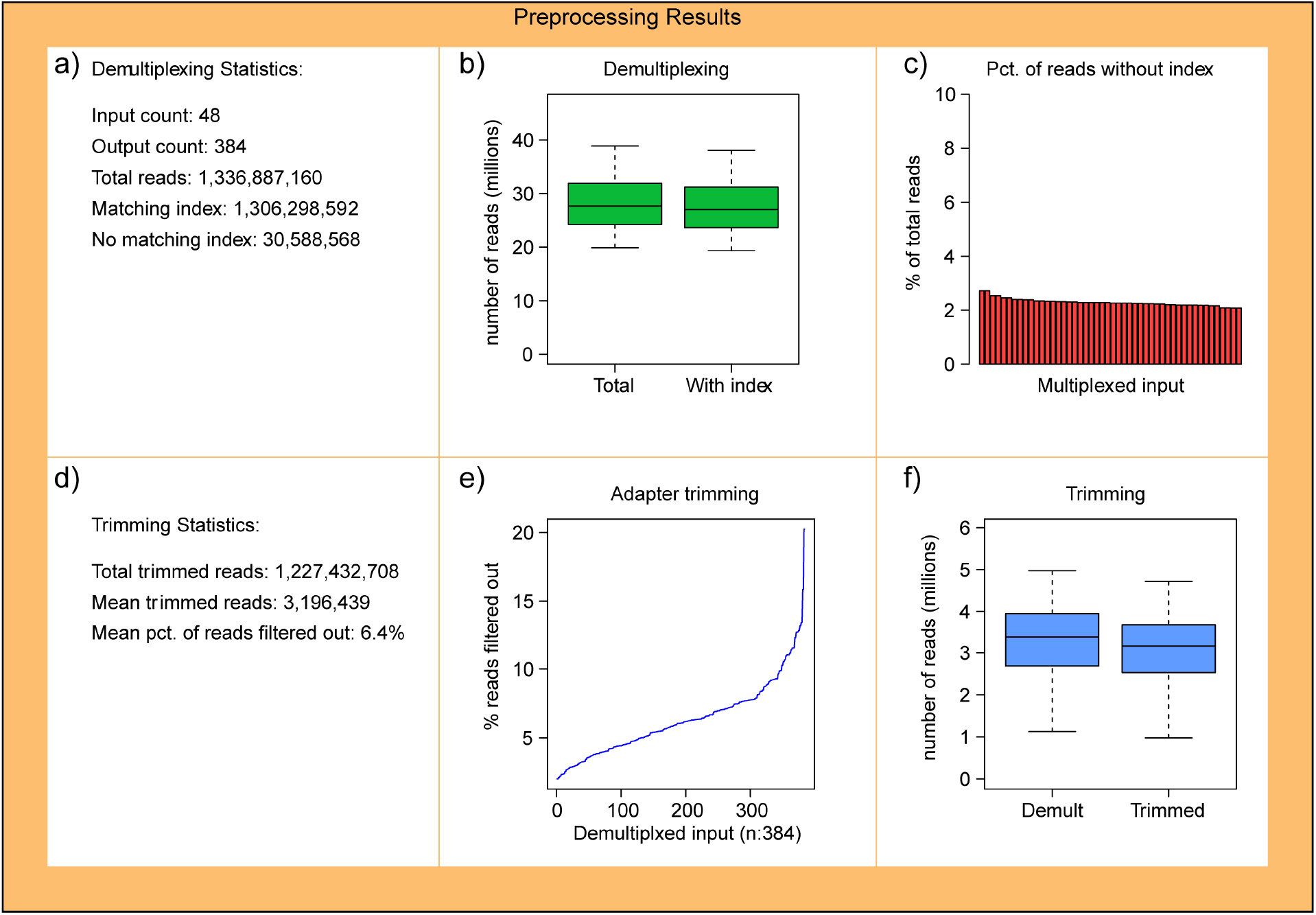
Example output of read preprocessing module. **a)** Overall demultiplexing statistics. Input count: The number of multiplexed input, which may correspond to lanes and wells in the sequencing flowcell. Output count: The number of demultiplexed output. If the sample is sequenced by single-ended format, this is the total number of cells. Otherwise, this is twice the number of cells. Matching/no matching index: the total number of reads having a valid or invalid index. **b)** Distribution of the total number of reads per input (left) and output (right). **c)** The percentage of reads missing a valid index per input, sorted from highest to lowest. **d)** Overall trimming statistics showing the total number of trimmed reads, mean number of trimmed reads per demultiplexed input, and mean percentage of reads filtered out per input. **e)** The distribution of the percentage of removed reads per input sorted from lowest to highest. **f)** The distribution of read counts per input before (left) and after (right) trimming.

SINBAD generates detailed statistics and graphical plots based on the analyses performed by each module. Although the pipeline can be used end-to-end, since it works with a wide range of standard data formats (Methods), it can also be used for specific data processing/analysis tasks only, such as methylation calling on aligned data or generating methylation matrix using cytosine calls. In either case, the outputs generated by SINBAD can be used directly by other downstream computational tools for single-cell genomics (Methods).

### Preprocessing

Unlike existing methods that assume the existence of a gene activity matrix, SINBAD starts with raw, multiplexed sequencing reads. SINBAD demultiplexes the raw reads using cell barcode sequence information, which is technology dependent. The indexed reads, which are defined as those that match the given indices, are generated for each individual cell as the output. Statistics such as the number of total reads and usable reads are summarized and presented as the quality metrics for demultiplexing (Fig. 1). The percentage of reads without index is also reported, to help identify any potential technical issues related to library preparation or sequencing (Fig. 1).

Since unmethylated cytosines are converted to thymines as a result of bisulfite treatment in BS-Seq experiments, they can map to both nucleotides (C and T), thus reducing sequencing complexity. This leads to a low alignment rate in methylome data. Untrimmed sequencing adapters can cause further reduction in alignment rate for methylation sequencing, which is already lower than other single-cell sequencing technologies, such as scRNA-Seq or scATAC-Seq. To address this issue, SINBAD implements the adapter trimming step following the demultiplexing steps. In the case of paired-end sequencing, different trimming settings for the left and right reads are supported, adding extra flexibility (Fig. 1).

### Mapping and Filtering

Read alignment is performed for each cell, and the distribution of alignment rates is plotted as a quality metric (Fig. 3). The aligned reads are filtered in multiple steps. First, the alignments with a MAPQ quality score below the threshold (an adjustable parameter) are filtered out. Next, clonal reads (due to PCR duplicates) and mapped reads that failed bisulfite conversion, which can be identified by the existence of consecutive methylated non-CpG sites in the read, are removed. If a spike-in control (such as lambda phage DNA) is used to measure bisulfite conversion rate, reads mapped to the target genome are separated from the control reads as the final clean reads to be used for the remaining data processing steps for each cell. The number of reads is recorded per cell, and the distributions are displayed for quality control (Fig. 3). If the input data consists of paired-end reads that are processed separately, as in the snmC-Seq protocol, left and right reads are merged in this step.

**Figure 3.**
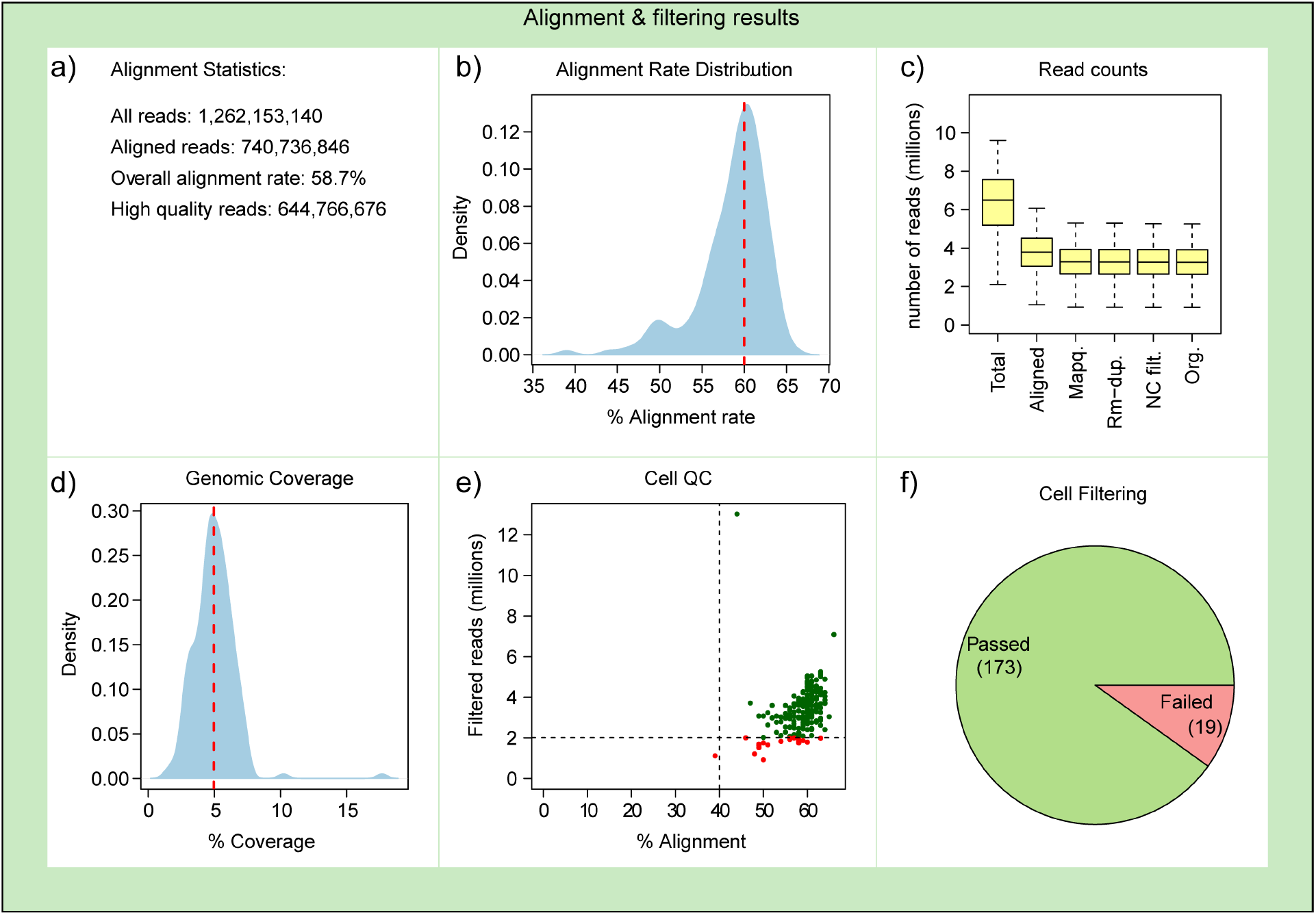
Example output of mapping and filtering module. **a)** Overall alignment statistics showing the total number of input reads for the sample, number of reads mapped to the reference genome, percentage of mapped reads among all input reads, and total number of aligned reads after filtering steps. **b)** Distribution of alignment rates per input. Dashed red line shows the median value. **c)** Distribution of total number of input reads, aligned to the genome, filtered by mapping quality, after removing clonal reads, after removing reads with failed bisulfite conversions, and after removing spike-in control reads. **d)** Distribution of the percentage of the genome covered by at least one read per cell. **e)** Scatter plot showing alignment rate (x axis) and total number of filtered reads (y axis) for each cell as the quality control (QC) metrics. The vertical and horizontal lines combined form the cell filtering cutoff. **f)** Pie chart showing the number of cells that passed QC thresholds and those that are discarded because of low quality.

One of the challenges for single-cell DNA methylome sequencing is to have sufficient coverage of the genome. Unlike RNA-Seq protocols, where the reads are highly enriched in coding regions, and ATAC-Seq protocols, where the fragments are from accessible chromatin regions, whole genome bisulfite sequencing data is scattered across the genome. As a result, even very deep sequencing experiments cannot reach >10% genomic coverage (Luo et al. 2017). Therefore, the coverage rate, if not the most definitive quality metric, is an important measure that can reflect the power of downstream analysis. Hence, SINBAD computes the genomic coverage rate, defined as the percentage of the genome that is covered by at least one read per cell (Fig. 3d).

Cells with low-quality data can cause undesired consequences in downstream analyses, such as clustering. Low mapping rates imply potential contamination, and a low number of aligned reads can limit robust analysis. We use two adjustable metrics, mapping rate and number of filtered reads, to exclude low-quality cells in SINBAD (Fig. 3e, 3f).

### Methylation calling

Using reads retained after the mapping and QC module, SINBAD calls methylated cytosines for cells passed QC. It calls methylated and unmethylated cytosines separately for the target genome and spike-in control if used in the experiment. Since the spike-in DNA is unmethylated, all cytosines are expected to be converted to thymines by bisulfite treatment in the ideal case. In this sense, the percentage of unmethylated cytosines to all cytosine calls in the spike-in control defines the conversion rate, which is one of the primary metrics used to evaluate the success of the experiment. SINBAD computes the conversion rate for each individual cell, stratified by the type of sequence context (CpG, CHG, CHH), and plots the overall statistics (Fig. 4a).

**Figure 4.**
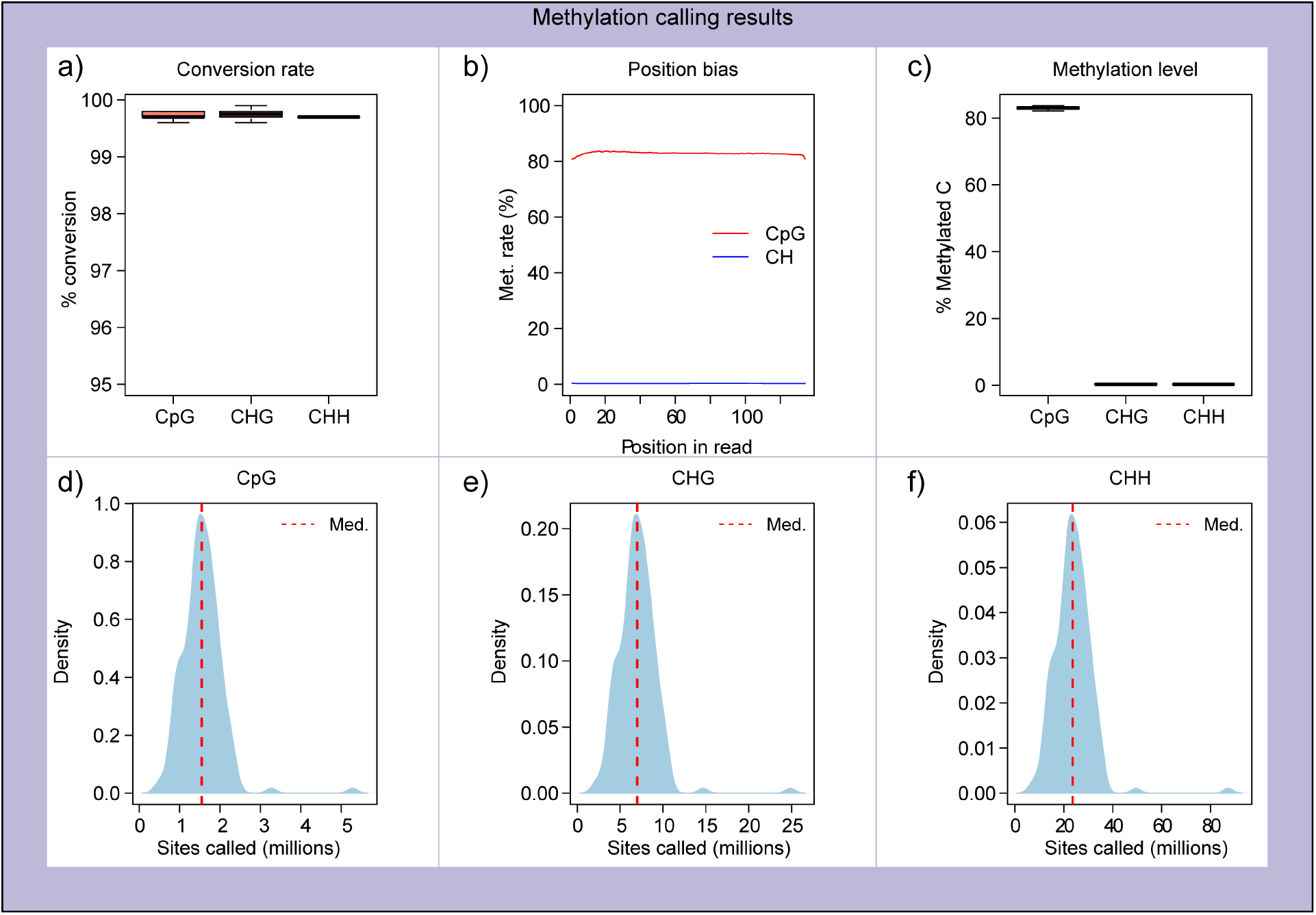
Example output of methylation module. **a)** Bisulfite conversion rate for CpG and non-CpG methylation, calculated using spike-in control DNA. **b)** Line plot showing positional methylation rate bias for cytosine sites in the read. **c)** Boxplots showing the methylation rate distributions for CpG and non-CpG sites. **d)** Distribution for the number of cytosine calls per cell for CpG sites **e)** CHG sites and **f)** CHH sites.

In addition to the overall bisulfite conversion rate in the data, the dependency of the conversion rate on the relative position of the cytosine in a read is another quality metric. Ideally, the conversion efficiency is expected to be independent of the relative position, and as a result, the methylation rates for both CpG and non-CpG sites are expected to be constant across the positions in the sequence. The position bias is calculated by SINBAD for both types of cytosine sites by combining all cells (Fig. 4b). Fluctuations in this metric indicate potential positional bias in the conversion rate. If fluctuations are located at the ends of the reads additional trimming may be needed.

Since DNA methylation predominantly occurs at CpG sites (Jang et al. 2017), CpG methylation rates are much higher than non-CpG methylation rates. Depending on the tissue or cell type under investigation, CpG methylation rate typically ranges between 60% and 80% for animal cells (Singer 2019). However, non-CpG rates are expected to be close to zero in animal cells, except for neurons, which can have as much as 5% methylation rate for non-CpG sites (Lee, Park, and Nakai 2017). Hence, non-CpG methylation rates can serve as another quality metric, as abnormally high methylation levels at such sites suggest potential technical issues, such as failed bisulfite conversion during sample processing. SINBAD computes the overall methylation rates on a per-cell basis for all three types of cytosine sites (Fig. 4 d,e,f), providing a preliminary technical and biological perspective of bisulfite conversion. The distribution of the number of cytosine calls is uneven throughout the genome due to regions with a high density of this nucleotide, such as CpG islands. Hence, the number of cytosine calls may not necessarily be the same as the overall genomic coverage of the aligned reads. To define the specific power of the generated data in terms of the cytosine sites covered, SINBAD computes the total number of cytosine calls stratified into three categories of cytosine sites (Fig. 4d, e, f).

### Quantification of methylation levels

Once the methylation sites are called, SINBAD quantifies the methylation levels for user-defined genomic regions for each cell. Due to the nature of DNA methylation, quantification of methylation levels requires extra flexibility compared to gene expression and chromatin accessibility data. For gene expression data, the genomic regions of interest are coding sequences. For chromatin accessibility data, the regions of interest are open chromatin regions defined by peaks.

For DNA methylation data, multiple types of genomic regions can be of interest, depending on the problem at hand and tissue/cell types. These could include the gene body, promoters, enhancers, insulators and other functional DNA elements. Finally, for single-cell data, dividing the genome into fixed-size bins can help to profile the heterogeneity present in the cell population. This binning approach can also assist in comparative analysis of methylation patterns across cells and samples in an unbiased manner.

SINBAD addresses the need for flexibility in methylation quantification by allowing user-defined genomic regions. Several sets of annotated genomic regions for human and mouse are included by default (Methods). Additionally, any user-defined set of regions can be processed. Given a set of genomic regions, SINBAD can quantify both CpG and non-CpG methylation levels for the set, generating a region-by-cell matrix as the output. These matrices can be used with any commonly used single-cell data analysis tool, such as Seurat, Monocle (Qiu et al. 2017) and SCANPY (Wolf, Angerer, and Theis 2018) for additional downstream analyses.

Before a methylation matrix can be used for downstream analysis, an additional processing step is required, which is unique to DNA methylation data. For gene expression and chromatin accessibility data, the number of normalized reads directly reflect gene expression level and chromatin accessibility. For DNA methylation data, the ratio of methylated cytosines to all cytosine calls constitutes the signal. Hence, the lack of cytosine calls for a region simply means missing information and does not necessarily mean lack of methylation, which must be handled before downstream analysis.

Due to the low genomic coverage of single-cell bisulfite sequencing data, short genomic regions likely lack sufficient numbers of cytosines to make a reliable estimate of the methylation level for the region, resulting in many missing values in the methylation matrix. Many dimensionality reduction methods commonly used in single-cell genomics are not compatible with missing values in the input matrix, hindering their utility for methylome data. We implemented a simple imputation technique in our pipeline (Methods) to support downstream analysis by using the population mean to replace missing values. More sophisticated imputation methods have been developed (Kapourani and Sanguinetti 2019; Uzun, Wu, and Tan 2020). The methylation calls generated by SINBAD can be used as the input to such tools, if needed.

### Downstream analysis

Two types of downstream analyses are implemented in SINBAD (Fig. 1). First, it performs dimensionality reduction using Principal Component Analysis (PCA) and Uniform Manifold Approximation and Projection (UMAP) on the methylation matrix generated by the quantification module. Next, cell clusters are identified in the low dimensional space. For cell type annotation, a set of marker genes can be provided for plotting their methylation levels, either on the UMAP or as a violin plot showing the distribution of methylation levels for the cells for each cluster.

As an unbiased analysis, SINBAD supports differential methylation analysis to identify the genes or functional DNA elements that have significantly higher or lower methylation levels among the clusters. The feature types to be investigated can be the same as the one that is used for the dimensionality reduction or any other feature types quantified earlier. The results can be used to assist cell type identification as well as discovery of novel genes and functional DNA elements associated with differential methylation.

### Case study

As a case study for demonstrating the utility of SINBAD, we obtained single-cell DNA methylation data for human frontal cortex generated using the snmC-Seq protocol (Luo et al. 2017). In this study, the authors processed the methylation data using in-house scripts and identified two main cell clusters that correspond to excitatory and inhibitory neuron subtypes, based on *non-CpG* methylation of known marker genes. Using SINBAD, we computed the *CpG* methylation levels across the genome using 100 kb genomic bins and performed dimensionality reduction. We were able to identify two cell clusters in the sample. The cluster separation was clearly observed when methylation levels of the genomic bins or gene bodies were used as the input. However, the cluster separation was poor when promoter methylation levels were used as the input, highlighting the need for flexible choice of genomic regions to be analyzed (Fig. 5a, Supp. Fig. 1).

**Figure 5.**
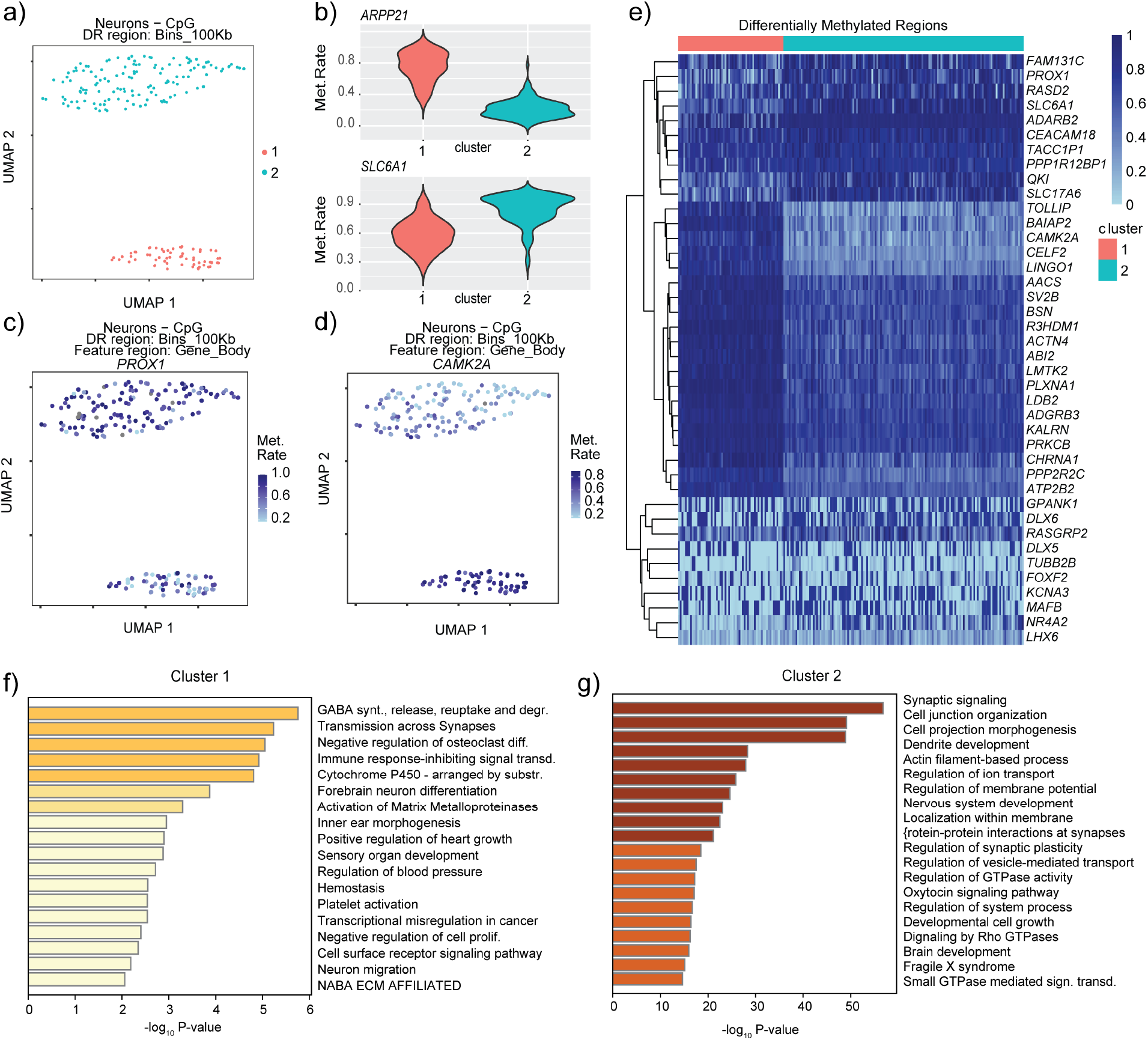
Case study of human frontal cortex methylome dataset. **a)** UMAP dimensionality reduction and clustering of cell populations. **b)** Violin plots showing the CpG methylation level of the gene body for the excitatory neuron marker *ARPP21* and the inhibitory neuron marker *SLC6A1*. **c)** UMAP showing the methylation levels of the gene body for the inhibitory neuron marker gene *PROX1* and **d)** the excitatory neuron marker gene *CAMK2A*. **e)** Heatmap for the differentially methylated genes between the two clusters. **f)** Enriched GO terms in the significantly demethylated genes in cluster 1 and **g)** cluster 2.

Profiling the methylation levels along the gene bodies and performing differentially methylated region (DMR) analysis between the two clusters revealed a consistent pattern of differential methylation of excitatory and inhibitory neuron marker genes between the clusters. Consistent with the results based on analyzing non-CpG methylation in the original study, the inhibitory neuron marker genes, including *SLC6A1*, *PROX1*, and *ADARB2*, were demethylated in one cluster (Fig. 5b, e), whereas the excitatory neuron marker genes, such as *ARPP21*, *BAIAP2*, and *CAMK2A*, were demethylated in the other cluster (Fig 5c, d, e) (Luo et al. 2017; Kang, Park, and Kim 2016).

In addition to known marker genes of the two neuronal subtypes, DMR analysis revealed many additional differentially methylated genes that have a role in neuronal development and function, including *RASD2*, *SLC17A6*, and *LINGO1*. Gene ontology analysis for the significantly demethylated genes in the cell clusters revealed terms that are associated with subtypes of neurons. The top enriched term for the demethylated genes in the inhibitory cluster was the life cycle of Gamma-aminobutyric acid (GABA), which is the primary inhibitory neurotransmitter (Jewett and Sharma 2018). The enriched terms for the significantly demethylated genes in the other cluster included those related to excitatory neurons, such as development of dendrites, which excitatory neurons typically have hundreds to thousands of (Miles et al. 1996; Kennedy 2016).

### Conclusion

SINBAD addresses a critical need for interoperable and efficient tools for single-cell DNA methylome data analysis. It provides a flexible framework for implementing various analysis tasks of single-cell DNA methylome data including preprocessing, quality control, methylation calling and quantification and additional downstream analyses. It generates both graphical and text reports on QC metrics and analysis results. By providing a standardized analysis framework, SINBAD facilitates the reproducibility of single-cell DNA methylation data analysis. Application of SINBAD on human frontal cortex dataset demonstrated its effectiveness in cell type annotation and identification of differentially methylated genes/regions.

Although the current version of SINBAD already provides a comprehensive set of analysis tools, additional tools and analyses can be incorporated in the future. For example, several alternative dimensionality reduction methods can be added, including tSNE (van der Maaten 2008), PHATE (Moon et al. 2019) or densMAP (Narayan, Berger, and Cho 2021). In addition, more sophisticated imputation methods such as Melissa ((Kapourani and Sanguinetti 2019) and CaMelia (Tang et al. 2021) can be added.

## Software availability

SINBAD is implemented as an open source software in R programming language and is publicly available at github (https://github.com/tanlabcode/SINBAD.1.0) under the MIT License.

## Acknowledgement

We thank Dr. Hao Wu for helpful discussion about the design of SINBAD. This work was supported by National Institutes of Health of United States of America grant CA233285 (to K.T.), a grant from the Leona M. and Harry B Helmsley Charitable Trust (2008-04062 to K.T.), and a grant from the Alex’s Lemonade Stand Foundation (to K.T.).

## Methods

### Read Preprocessing

The first step of the data processing pipeline is demultiplexing the sequencing reads in fastq format. To reduce storage needs, SINBAD can handle the data in compressed format. The demultiplexing script *demultiplex_fastq.pl* can be called in R. The index length, which is sequencing protocol dependent, can be set by the user with the demux_index_length parameter and is used to demarcate the index sequence to demultiplex the data into multiple cells.

For some protocols, such as snmC-Seq and snmC-Seq2, the index is only present in the left reads in paired-end data. Our pipeline supports such protocols by generating intermediate right reads with an index prior to demultiplexing, which are removed after demultiplexing. SINBAD uses Cutadapt (Martin 2011) or Trimmomatic (Bolger, Lohse, and Usadel 2014) for trimming sequencing adapters.

### Read alignment

Trimmed reads are mapped to the reference genome using Bismark (Krueger and Andrews 2011) and alignment files in bam format are generated as the output. If spike-in control such as lambda phage DNA is used to measure the bisulfite conversion rate, the phage genomic sequence should be added to the reference genome sequence file as a separate chromosome. By default, we provide the hg38 and mm10 reference genome sequence (including lambda phage) with SINBAD.

Low quality alignments are filtered out based on the mapq_threshold parameter set by the user and clonal reads are removed using the samtools rmdup utility (Li et al. 2009; Li 2011). The aligned reads with failed bisulfite conversion, which is indicated by the presence of three consecutive non-CpG methylation events, are also removed. If a spike-in control is used, the reads mapped to the control sequence are separated from the target genome for downstream analyses, and the remaining reads are sorted and indexed using the samtools. The number of aligned reads is calculated using the countBam function, and the coverage rates of the mapped reads per cell are calculated using the pileup function of the Rsamtools package (Morgan et al. 2016).

Following the alignment step, low quality cells, which are defined as having alignment rate below the minimum alignment rate (set by the alignment_rate_threshold parameter) and aligned filtered reads below the threshold (set by the total_minimum_filtered_read_count parameter), are marked in the data. Only high quality cells passing these filters are used in downstream analyses.

### Methylation calling

SINBAD uses Bismark Methylation Extractor (Krueger and Andrews 2011) to call methylated cytosines, using the aligned and filtered reads as the input. This step is executed separately for the spike-in control (if used) which in turn is used to compute the bisulfite conversion rate distribution for the cells. Methylation calling results are generated for both CpG and non-CpG sites and all are converted into bed format to facilitate its usage by various external tools. Methylation calls in bed format are generated as output as a result of this step.

### Quantification of methylation level

We designed SINBAD to accept any set of genomic regions in bed format, such as gene bodies, promoters, or genomic bins of fixed sizes. By default, we provide several sets of predefined genomic regions with the software, including gene bodies, promoters and genomic bins of 10Kb and 100Kb size.

We determine the number of methylated and unmethylated cytosines in a given genomic region using GenomicRanges (Lawrence et al. 2013) and then calculate the average methylation rate per cell and per region by dividing the total number of methylated cytosines by all cytosine called. Due to the sparse nature of DNA methylation data, many cells are expected to have an insufficient number of cytosine calls, set by the min_call_count_threshold_for_region parameter, to estimate the methylation level for the cells. We impute the methylation level for a given region and a cell by using the mean rate of the same region based on cells with a sufficient number of cytosine calls. The generated methylation matrices are saved in serialized R data object (RDS) format, which can be used as the input to commonly used single-cell analysis tools such as Seurat (Hao et al. 2021) and Monocle (Qiu et al. 2017).

### Downstream analysis

SINBAD performs dimensionality reduction on the methylation matrix generated by the quantification module. First, using the top 2,000 regions with largest variance in the data as the input, principal components of the methylation matrix are identified with the prcomp function. Next, low-dimensional representation of the data is generated using UMAP (McInnes et al. 2018) and the top principal components explaining the largest variance in the data. In the lowdimensional feature space, density clustering is employed to identify cell populations in the sample (Rodriguez and Laio 2014). Methylation rates for the user defined marker genes or regions can be visualized on the UMAP and violin plots.

Differential methylation analysis is performed for each genomic region between two cell clusters using t-test or Wilcoxon rank sum test. Only cells having sufficient number of cytosine calls to estimate the methylation level for that region are used. P-values are corrected for multiple testing with the Benjamini-Hochberg method. A heatmap showing the expression levels of significantly differentially methylated regions is generated using the pheatmap package (Kolde and Others 2012).

### Processing of frontal cortex dataset

We processed the snmC-Seq dataset of human frontal cortex (McInnes et al. 2018; Luo et al. 2017) as follows. Bismark cov files for the methylation calls were obtained from GEO (accession GSE97179). We quantified CpG methylation calls for each cell across gene bodies (from transcription start sites (TSS) to the end of 3’ UTR), promoters (upstream and downstream 2Kb of TSS) and genomic bins (100Kb and 10Kb) for 200 cells. UMAP dimensionality reduction was applied for each of the feature matrices above on the top 10 PCA components responsible for the largest variance in the data. For the regions without sufficient number of CpG calls for methylation level estimate, we used the mean methylation rate of the entire population for that region. Density clustering was employed to identify cell clusters, and t-test was used to identify the differentially methylated genes. We then inspected methylation levels of excitatory and inhibitory neuron marker genes and performed differential methylation analysis with SINBAD. GO terms enriched in differentially methylated genes across the clusters (adjusted p < 0.01) were computed using metascape (Zhou et al. 2019).

### Parallelization

SINBAD is implemented with the R statistical programming language (Wirtschaftsuniversität Wien Department of Statistics and Mathematics 2008). To increase processing speed, we implemented parallel processing with multiple threads by using R doSNOW package (Analytics and Weston 2014) in SINBAD. For the demultiplexing step, the multiplexed inputs can be demultiplexed individually by a separate thread. In the same manner, all the remaining steps can be parallelized for the individual cells using isolated processes with user-specified number of threads, until dimensionality reduction. This flexibility allows the user to adjust the balance between memory and CPU allocation and running time. We noticed that Bismark alignment can terminate prematurely when the allocated memory becomes insufficient. This is remedied in SINBAD by additional alignment attempts for the cells for which the mapping thread fails.

