## Supplemental Material for "SINBAD: a flexible tool for single cell DNA methylation data"

**Supplementary Materials**

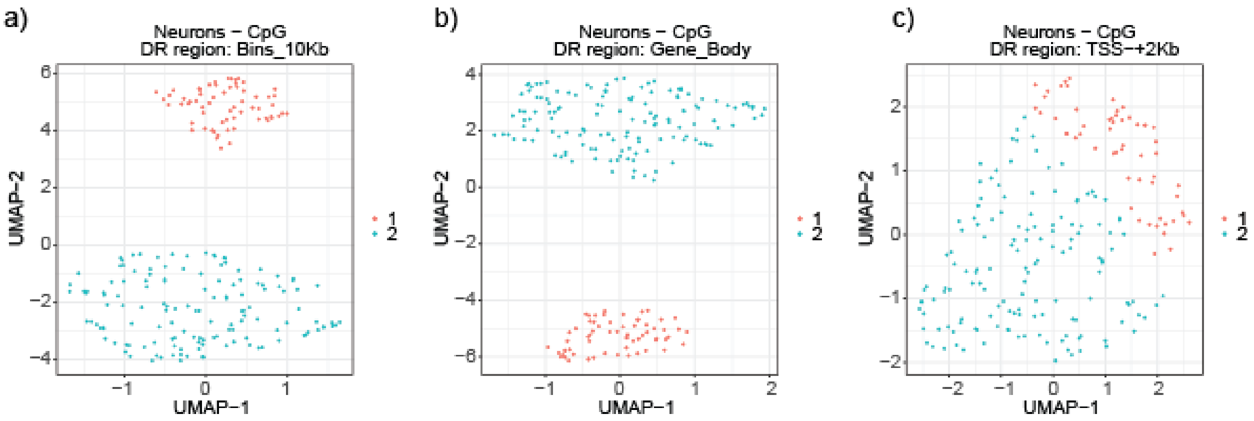


**Supplementary Figure 1. Dimensionality reduction and clustering using CpG methylation for frontal cortex dataset (GSE97179) generated with snmC-Seq protocol. a)** Using 10 Kb length genomic bins as the input matrix **b)** Using gene body methylation **c)** Using promoters, defined as the region extending along 2 Kb upstream and 2 Kb downstream of transcription start sites. DR: Dimensionality reduction
